## Supplemental figures and tables for "Local cortical desynchronization and pupil-linked arousal differentially shape brain states for optimal sensory performance"

Leonhard Waschke*, Sarah Tune, & Jonas Obleser*
University of Lübeck, Germany

*Author correspondence:
Leonhard Waschke, Jonas Obleser
Department of Psychology
University of Lübeck
Maria-Goeppert Straße 9a
23562 Lübeck


**Supplemental figures**

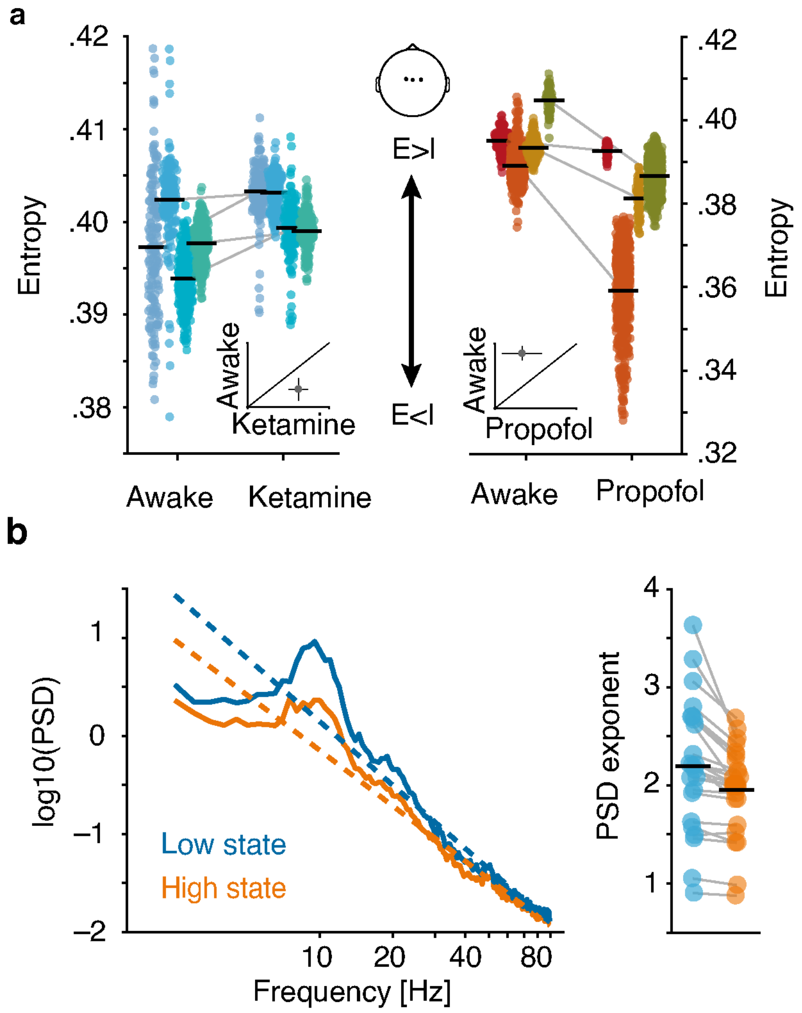


**Figure 2 supplement 1. Validation of EEG entropy as a marker of cortical excitation / inhibition balance**. (**a**) EEG entropy of four participants (averaged over three central channels) recorded in awake state and under ketamine anesthesia (left panel). Single dots represent snippets of data, black vertical lines depict the average within a participant. EEG entropy of four different participants recorded in awake state and under propofol anesthesia (right panel). Insets illustrate the grand-average (± SEMs) using a 45° plot. Excitatory activity is expected to be relatively increased during ketamine anesthesia and decreased after propofol infusion (central arrow). (**b**) Grand-average power spectral density of high and low entropy states (left panel) including illustrative fitted slopes. PSD exponents of single participants (right panel).

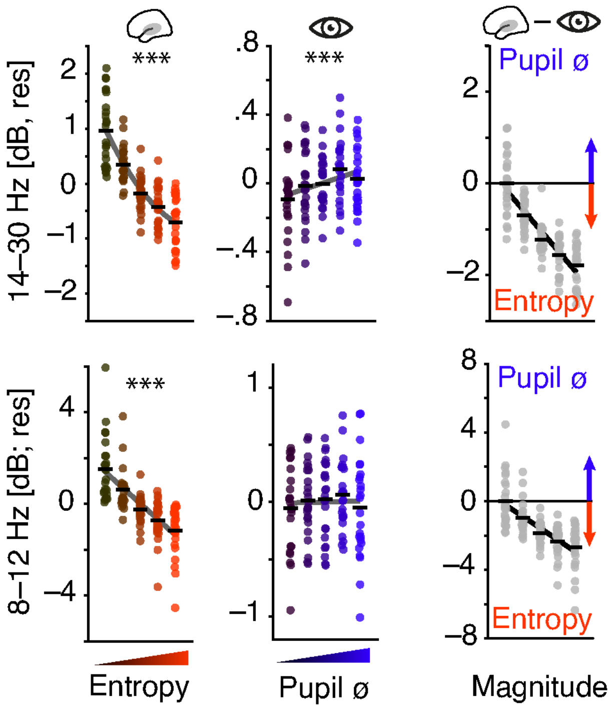


**Figure 3 supplement 1. Ongoing activity in the alpha and beta band as a function of EEG entropy and pupil size.** Pre-stimulus EEG entropy (red colours) and oscillatory power are negatively linked in the alpha band (8–12 Hz, lower panels) and slightly weaker in the beta band (14–30 Hz, upper panels). Pre-stimulus pupil size (blue colours) is positively related to beta but not alpha power.

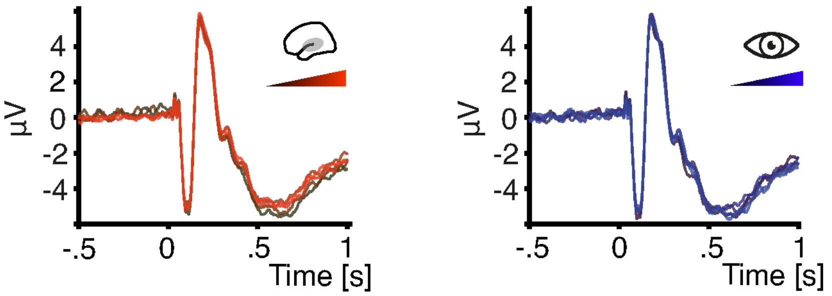


**Figure 4 supplement 1. Grand average ERPs for increasing pre-stimulus entropy and pupil dilation.** Grand average ERP time-courses over auditory cortical areas (spatial filter) for five bins of increasing pre-stimulus entropy (left) and pupil size (right). Baseline-corrected to 500 ms pre-stimulus.

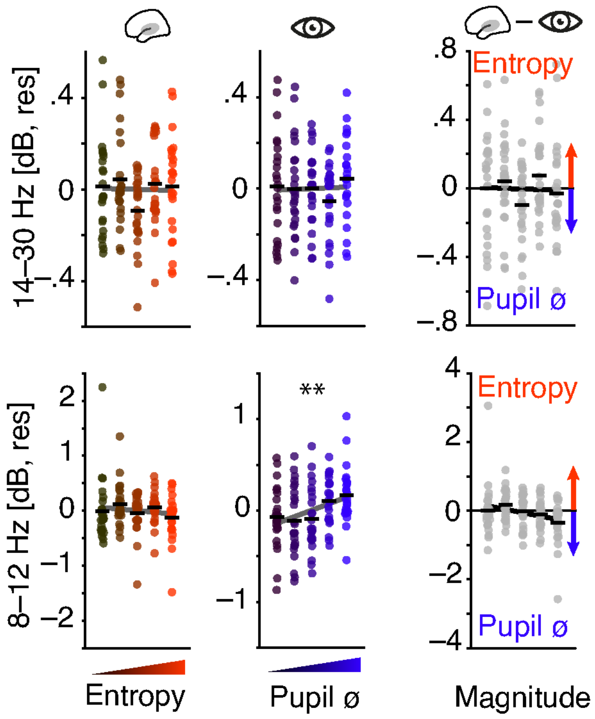


**Figure 4 supplement 2. Tone-related activity in the alpha and beta band as a function of pre-stimulus EEG entropy and pupil size.** Pre-stimulus EEG entropy (red colours) and sensory-evoked oscillatory power are not substantially linked in the alpha band (8–12 Hz, lower panels) or the beta band (14–30 Hz, upper panels). Pre-stimulus pupil size (blue colours) positively correlates with sensory-evoked alpha power.

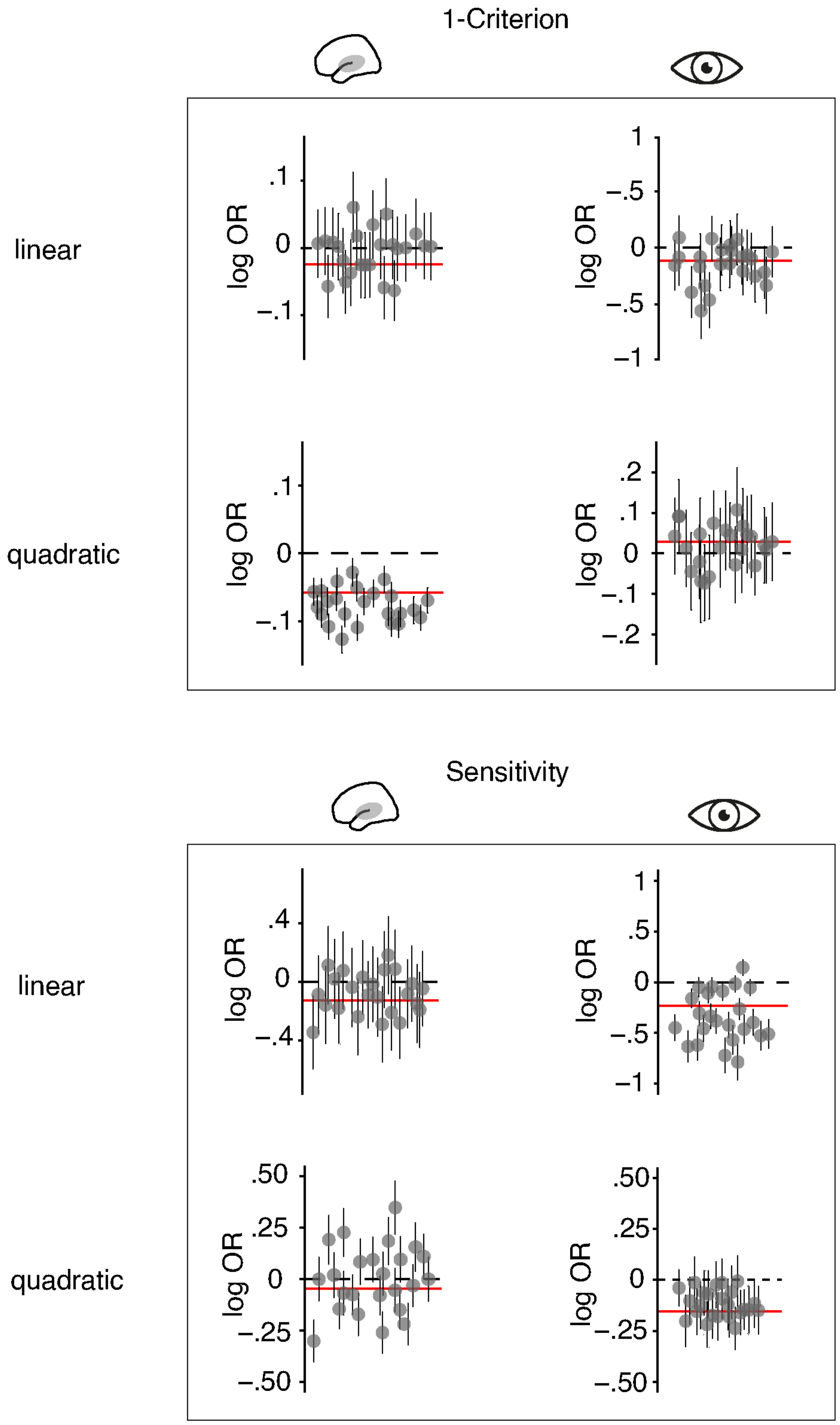


**Figure 5 supplement 1. Overview of fixed and random effects.** Effects of auditory cortical entropy (left) and pupil size (right) on 1-criterion (top) and sensitivity (bottom). Entropy either calculated based on EEG-signals from auditory (left) or visual cortical areas (right). Linear and quadratic effects shown. As above, grey dots represent single subject random slopes (including 95% CIs) and red lines fixed effects in log odds ratios (log OR). Model details in Table S6.

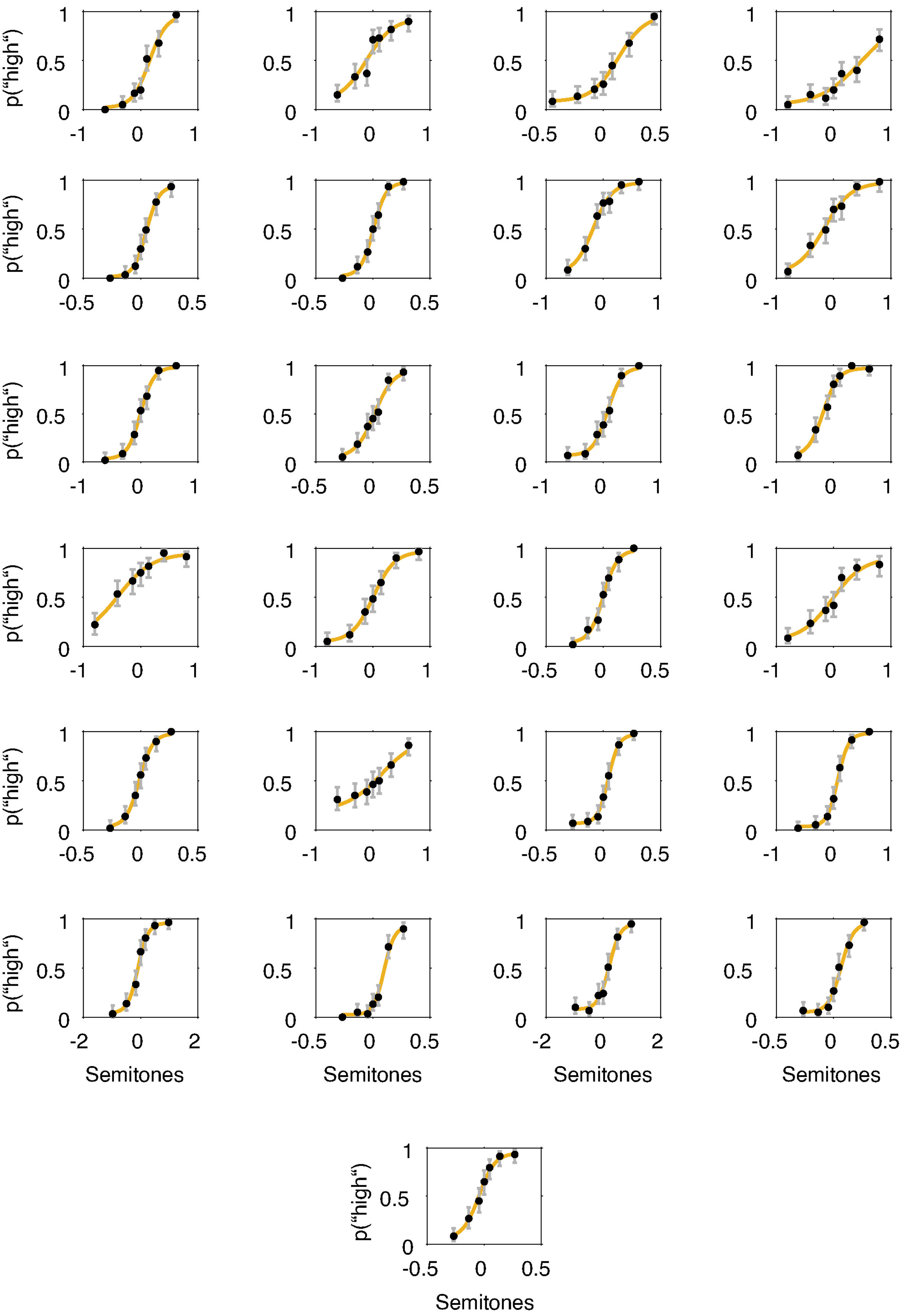


**Figure 5 supplement 2. Single-participant psychometric functions.** Average probability of “high” responses across seven levels of stimulus pitch (semitones relative to the median frequency) for all 25 participants. Black dots represent actual data, grey lines 95% confidence intervals, and yellow lines fitted psychometric functions. Note that the varying range of stimulus pitch stems from the initial participant-wise titration procedure.

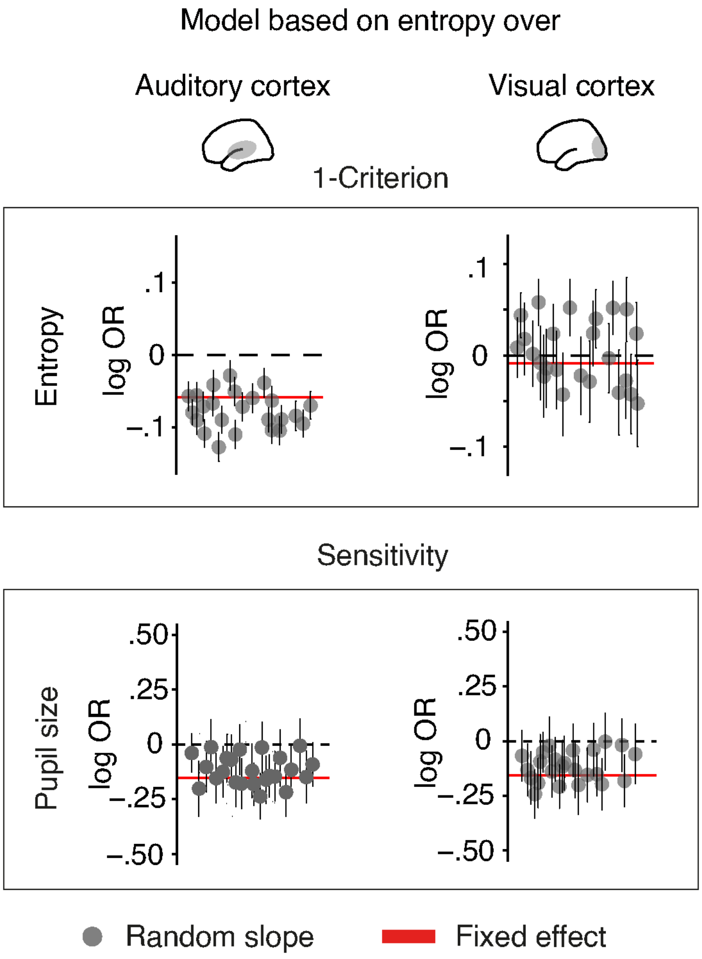


**Figure 6 supplement 1. Comparison of results from different brain–behaviour models.** Effects of entropy (top) and pupil size (bottom) on Criterion and sensitivity, respectively. Entropy either calculated based on EEG-signals from auditory (left) or visual cortical areas (right). Red lines represent fixed effects, grey dots single subject random slopes (including 95% CIs). Note that visual cortical entropy does not predict Criterion changes (top right) but the overall influence of pupil size on sensitivity remains unchanged by changing the cortical origin of entropy (bottom). Model details in Table S6 and S8.

### Supplementary tables

**Table S1: Brain-brain model predicting pre-stimulus low-frequency power**

| | **Pre-stimulus low frequency power** | | | | |
|---|---|---|---|---|---|
| *Predictors* | *Estimates* | *std. Error* | *CI* | *t-value* | *p* |
| Intercept | -0.041 | 0.038 | -0.116 – 0.033 | -1.089 | 0.2760 |
| **Entropy (linear)** | **-0.179** | **0.011** | **-0.200 – -0.158** | **-16.570** | **<0.001** |
| **Entropy (quadratic)** | **0.030** | **0.009** | **0.012 – 0.048** | **3.299** | **0.0010** |
| Entropy baseline | 0.045 | 0.013 | 0.020 – 0.070 | 3.519 | 0.0004 |
| **Pupil size (linear)** | **-0.041** | **0.010** | **-0.062 – -0.021** | **-3.923** | **0.0001** |
| **Pupil size (quadratic)** | **0.016** | **0.006** | **0.003 – 0.028** | **2.504** | **0.0123** |
| Entropy (linear) x Baseline | -0.000 | 0.001 | -0.003 – 0.003 | -0.071 | 0.9434 |
| Entropy(quadratic) x Baseline | -0.013 | 0.010 | -0.033 – 0.007 | -1.249 | 0.2115 |
| Participant | 0.007 | 0.007 | -0.006 – 0.020 | 1.014 | 0.3106 |
| Observations | 9831 | | | | |
| $R^2$ / adjusted $R^2$ | 0.033 / 0.032 | | | | |

**Table S2: Brain-brain model predicting pre-stimulus alpha power**

| | Pre-stimulus alpha power | | | | |
|---|---|---|---|---|---|
| *Predictors* | *Estimates* | *std. Error* | *CI* | *t-value* | *p* |
| Intercept | -0.047 | 0.037 | -0.120 – 0.026 | -1.268 | 0.2049 |
| **Entropy (linear)** | **-0.291** | **0.011** | **-0.311 – -0.270** | **-27.575** | **<0.001** |
| **Entropy (quadratic)** | **0.057** | **0.009** | **0.039 – 0.074** | **6.372** | **<0.001** |
| Entropy baseline | 0.077 | 0.012 | 0.052 – 0.101 | 6.204 | <0.001 |
| Pupil size (linear) | 0.009 | 0.010 | -0.010 – 0.029 | 0.930 | 0.3524 |
| Pupil size (quadratic) | -0.004 | 0.006 | -0.016 – 0.009 | -0.566 | 0.5715 |
| Entropy (linear) x Baseline | -0.000 | 0.001 | -0.003 – 0.003 | -0.097 | 0.9230 |
| Entropy(quadratic) x Baseline | -0.019 | 0.010 | -0.038 – 0.001 | -1.880 | 0.0601 |
| Participant | 0.011 | 0.007 | -0.002 – 0.024 | 1.651 | 0.0988 |
| Observations | 9831 | | | | |
| $R^2$ / adjusted $R^2$ | 0.082 / 0.082 | | | | |

**Table S3: Brain-brain model predicting pre-stimulus beta power**

| | Pre-stimulus beta power | | | | |
|---|---|---|---|---|---|
| *Predictors* | *Estimates* | *std. Error* | *CI* | *t-value* | *p* |
| Intercept | -0.062 | 0.037 | -0.134 – 0.010 | -1.699 | 0.0893 |
| **Entropy (linear)** | **-0.316** | **0.010** | **-0.336 – -0.296** | **-30.329** | **<0.001** |
| **Entropy (quadratic)** | **0.074** | **0.009** | **0.056 – 0.091** | **8.336** | **<0.001** |
| Entropy baseline | 0.106 | 0.012 | 0.082 – 0.130 | 8.701 | <0.001 |
| **Pupil size (linear)** | **0.043** | **0.010** | **0.023 – 0.063** | **4.267** | **<0.001** |
| Pupil size (quadratic) | -0.004 | 0.006 | -0.016 – 0.008 | -0.582 | 0.5608 |
| Entropy (linear) x Baseline | -0.000 | 0.001 | -0.003 – 0.002 | -0.106 | 0.9158 |
| Entropy(quadratic) x Baseline | -0.026 | 0.010 | -0.045 – -0.006 | -2.603 | 0.0093 |
| Participant | 0.014 | 0.007 | 0.001 – 0.027 | 2.128 | 0.0333 |
| Observations | 9831 | | | | |
| $R^2$ / adjusted $R^2$ | 0.102 / 0.102 | | | | |

**Table S4: Brain-brain model predicting pre-stimulus gamma power**

| | Pre-stimulus gamma power | | | | |
|---|---|---|---|---|---|
| *Predictors* | *Estimates* | *std. Error* | *CI* | *t-value* | *p* |
| Intercept | -0.054 | 0.038 | -0.129 – 0.020 | -1.426 | 0.1539 |
| **Entropy (linear)** | **-0.179** | **0.011** | **-0.200 – -0.157** | **-16.543** | **<0.001** |
| **Entropy (quadratic)** | **0.061** | **0.009** | **0.043 – 0.079** | **6.655** | **<0.001** |
| Entropy baseline | 0.049 | 0.013 | 0.024 – 0.074 | 3.842 | 0.0001 |
| Pupil size (linear) | -0.013 | 0.010 | -0.034 – 0.007 | -1.275 | 0.2022 |
| Pupil size (quadratic) | 0.001 | 0.006 | -0.012 – 0.013 | 0.127 | 0.8992 |
| Entropy (linear) x Baseline | -0.000 | 0.001 | -0.003 – 0.003 | -0.066 | 0.9474 |
| Entropy(quadratic) x Baseline | -0.022 | 0.010 | -0.042 – -0.002 | -2.178 | 0.0294 |
| Participant | 0.001 | 0.007 | -0.012 – 0.014 | 0.148 | 0.8822 |
| Observations | 9831 | | | | |
| $R^2$ / adjusted $R^2$ | 0.035 / 0.034 | | | | |

**Table S5: Brain-brain model predicting post-stimulus low-frequency power**

| | Stimulus-evoked low frequency power | | | | |
|---|---|---|---|---|---|
| *Predictors* | *Estimates* | *std. Error* | *CI* | *t-value* | *p* |
| Intercept | 0.001 | 0.039 | -0.074 – 0.077 | 0.038 | 0.9694 |
| **Entropy (linear)** | **-0.026** | **0.011** | **-0.048 – -0.005** | **-2.387** | **0.0170** |
| Entropy (quadratic) | 0.001 | 0.009 | -0.017 – 0.019 | 0.107 | 0.9149 |
| Entropy baseline | 0.043 | 0.013 | 0.018 – 0.069 | 3.376 | 0.0007 |
| Pupil size (linear) | 0.015 | 0.011 | -0.006 – 0.035 | 1.369 | 0.1711 |
| Pupil size (quadratic) | 0.010 | 0.006 | -0.002 – 0.023 | 1.604 | 0.1088 |
| Entropy (linear) x Baseline | -0.000 | 0.001 | -0.003 – 0.003 | -0.159 | 0.8740 |
| Entropy(quadratic) x Baseline | -0.038 | 0.010 | -0.058 – -0.018 | -3.660 | 0.0003 |
| Participant | 0.014 | 0.007 | 0.000 – 0.027 | 2.000 | 0.0456 |
| Observations | 9831 | | | | |
| $R^2$ / adjusted $R^2$ | 0.006 / 0.005 | | | | |

**Table S6: Brain-brain model predicting post-stimulus alpha power**

| | Post-stimulus alpha power | | | | |
|---|---|---|---|---|---|
| *Predictors* | *Estimates* | *std. Error* | *CI* | *t-value* | *p* |
| Intercept | 0.012 | 0.039 | -0.064 – 0.088 | 0.311 | 0.7558 |
| Entropy (linear) | -0.008 | 0.011 | -0.029 – 0.014 | -0.706 | 0.4801 |
| Entropy (quadratic) | -0.011 | 0.009 | -0.029 – 0.007 | -1.160 | 0.2461 |
| Entropy baseline | 0.043 | 0.013 | 0.018 – 0.069 | 3.374 | 0.0007 |
| **Pupil size (linear)** | **0.033** | **0.011** | **0.013 – 0.054** | **3.141** | **0.0017** |
| Pupil size (quadratic) | -0.000 | 0.006 | -0.013 – 0.012 | -0.027 | 0.9785 |
| Entropy (linear) x Baseline | -0.000 | 0.001 | -0.003 – 0.003 | -0.042 | 0.9669 |
| Entropy(quadratic) x Baseline | -0.002 | 0.010 | -0.022 – 0.019 | -0.145 | 0.8847 |
| Participant | 0.007 | 0.007 | -0.007 – 0.020 | 0.947 | 0.3438 |
| Observations | 9831 | | | | |
| $R^2$ / adjusted $R^2$ | 0.004 / 0.003 | | | | |

**Table S7: Brain-brain model predicting post-stimulus beta power**

| | Post-stimulus beta power | | | | |
|---|---|---|---|---|---|
| *Predictors* | *Estimates* | *std. Error* | *CI* | *t-value* | *p* |
| Intercept | -0.014 | 0.039 | -0.089 – 0.062 | -0.349 | 0.7268 |
| Entropy (linear) | -0.006 | 0.011 | -0.027 – 0.016 | -0.516 | 0.6059 |
| Entropy (quadratic) | 0.001 | 0.009 | -0.017 – 0.020 | 0.150 | 0.8806 |
| Entropy baseline | 0.007 | 0.013 | -0.019 – 0.032 | 0.521 | 0.6026 |
| Pupil size (linear) | 0.007 | 0.011 | -0.014 – 0.028 | 0.656 | 0.5122 |
| Pupil size (quadratic) | 0.008 | 0.006 | -0.004 – 0.021 | 1.292 | 0.1963 |
| Entropy (linear) x Baseline | 0.000 | 0.001 | -0.003 – 0.003 | 0.026 | 0.9790 |
| Entropy(quadratic) x Baseline | 0.012 | 0.010 | -0.009 – 0.032 | 1.128 | 0.2594 |
| Participant | 0.002 | 0.007 | -0.012 – 0.015 | 0.225 | 0.8218 |
| Observations | 9831 | | | | |
| $R^2$ / adjusted $R^2$ | 0.001 / -0.000 | | | | |

**Table S8: Brain-brain model predicting post-stimulus gamma power**

| | **Post-stimulus gamma power** | | | | |
|---|---|---|---|---|---|
| *Predictors* | *Estimates* | *std. Error* | *CI* | *t-value* | *p* |
| Intercept | 0.012 | 0.039 | -0.064 – 0.088 | 0.313 | 0.7546 |
| **Entropy (linear)** | **0.040** | **0.011** | **0.018 – 0.061** | **3.598** | **0.0003** |
| Entropy (quadratic) | -0.007 | 0.009 | -0.026 – 0.011 | -0.783 | 0.4337 |
| Entropy baseline | 0.013 | 0.013 | -0.012 – 0.038 | 1.021 | 0.3074 |
| Pupil size (linear) | -0.016 | 0.011 | -0.037 – 0.004 | -1.539 | 0.1239 |
| Pupil size (quadratic) | 0.004 | 0.006 | -0.008 – 0.017 | 0.671 | 0.5023 |
| Entropy (linear) x Baseline | -0.000 | 0.001 | -0.003 – 0.003 | -0.069 | 0.9449 |
| Entropy(quadratic) x Baseline | -0.029 | 0.010 | -0.050 – -0.009 | -2.798 | 0.0051 |
| Participant | -0.003 | 0.007 | -0.017 – 0.010 | -0.458 | 0.6468 |
| Observations | 9831 | | | | |
| $R^2$ / adjusted $R^2$ | 0.003 / 0.002 | | | | |

**Table S9: Brain-brain model predicting post-stimulus ITC**

| | Post-stimulus low frequency ITC | | | | |
|---|---|---|---|---|---|
| *Predictors* | *Estimates* | *std. Error* | *CI* | *t-value* | *p* |
| Intercept | 0.035 | 0.039 | -0.041 – 0.110 | 0.897 | 0.3697 |
| **Entropy (linear)** | **0.052** | **0.011** | **0.031 – 0.074** | **4.759** | **<0.001** |
| **Entropy (quadratic)** | **-0.022** | **0.009** | **-0.040 – -0.004** | **-2.361** | **0.0182** |
| Entropy baseline | -0.016 | 0.013 | -0.042 – 0.009 | -1.276 | 0.2021 |
| Pupil size (linear) | 0.002 | 0.011 | -0.018 – 0.023 | 0.227 | 0.8207 |
| Pupil size (quadratic) | -0.005 | 0.006 | -0.017 – 0.008 | -0.720 | 0.4715 |
| Entropy (linear) x Baseline | -0.000 | 0.001 | -0.003 – 0.003 | -0.096 | 0.9239 |
| Entropy(quadratic) x Baseline | -0.026 | 0.010 | -0.046 – -0.005 | -2.473 | 0.0134 |
| Participant | 0.011 | 0.007 | -0.002 – 0.025 | 1.644 | 0.1002 |
| Observations | 9831 | | | | |
| $R^2$ / adjusted $R^2$ | 0.005 / 0.004 | | | | |

**Table S10: Brain-behavior model predicting decisions (high vs. low)**

| | Decision | | | | |
|---|---|---|---|---|---|
| *Predictors* | *Log-Odds* | *std. Error* | *CI* | *z-value* | *p* |
| Intercept | -0.057 | 0.148 | -0.347 – 0.233 | -0.384 | 0.701 |
| Pitch | 3.776 | 0.216 | 3.354 – 4.199 | 17.513 | <0.001 |
| Entropy (linear) | 0.024 | 0.030 | -0.034 – 0.082 | 0.801 | 0.423 |
| **Entropy (quadratic)** | **-0.059** | **0.026** | **-0.109 – -0.009** | **-2.311** | **0.021** |
| Baseline Entropy | 0.002 | 0.035 | -0.067 – 0.070 | 0.051 | 0.959 |
| **Pupil size (linear)** | **0.115** | **0.028** | **0.059 – 0.170** | **4.049** | **<0.001** |
| Pupil size (quadratic) | 0.028 | 0.016 | -0.004 – 0.060 | 1.722 | 0.085 |
| Trial number | 0.073 | 0.027 | 0.020 – 0.126 | 2.679 | 0.007 |
| Pitch x Entropy (linear) | -0.126 | 0.070 | -0.263 – 0.012 | -1.788 | 0.074 |
| Pitch x Entropy (quadratic) | -0.047 | 0.058 | -0.161 – 0.066 | -0.817 | 0.414 |
| Pitch x Baseline Entropy | -0.022 | 0.085 | -0.190 – 0.145 | -0.262 | 0.793 |
| Entropy (linear) x Baseline Entropy | 0.090 | 0.029 | 0.033 – 0.147 | 3.104 | 0.002 |
| Entropy (quadratic) x Baseline Entropy | 0.007 | 0.019 | -0.030 – 0.045 | 0.388 | 0.698 |
| **Pitch x Pupil size (linear)** | **-0.232** | **0.068** | **-0.365 – -0.098** | **-3.405** | **0.001** |
| **Pitch x Pupil size (quadratic)** | **-0.153** | **0.035** | **-0.222 – -0.085** | **-4.371** | **<0.001** |
| Pitch x Entropy (linear) x Baseline Entropy | 0.058 | 0.070 | -0.079 – 0.195 | 0.825 | 0.409 |
| Pitch x Entropy (quadratic) x Baseline Entropy | -0.002 | 0.045 | -0.091 – 0.087 | -0.040 | 0.968 |
| **Random Effects** | | | | | |
| $\sigma^2$ | 3.29 | | | | |
| $\tau_{00\ id}$ | 0.49 | | | | |
| $\tau_{11\ id.semitones}$ | 0.86 | | | | |
| $\rho_{01\ id}$ | -0.31 | | | | |
| Observations | 9831 | | | | |
| Marginal $R^2$ / Conditional $R^2$ | 0.542 / 0.632 | | | | |

**Table S11: Brain-behavior model predicting decisions (high vs. low)**

| | Decision | | | | |
|---|---|---|---|---|---|
| *Predictors* | *Log-Odds* | *std. Error* | *CI* | *z-value* | *p* |
| Intercept | -0.131 | 0.146 | -0.418 – 0.155 | -0.898 | 0.3694 |
| Pitch | 3.757 | 0.213 | 3.339 – 4.175 | 17.618 | <0.001 |
| Visual cortex entropy (linear) | 0.006 | 0.034 | -0.060 – 0.072 | 0.185 | 0.8529 |
| **Visual cortex entropy (quadratic)** | **0.009** | **0.021** | **-0.032 – 0.050** | **0.435** | **0.6633** |
| Baseline Entropy | -0.022 | 0.034 | -0.090 – 0.045 | -0.647 | 0.5179 |
| **Pupil size (linear)** | **0.112** | **0.029** | **0.056 – 0.168** | **3.925** | **0.0001** |
| Pupil size (quadratic) | 0.026 | 0.016 | -0.006 – 0.058 | 1.607 | 0.1082 |
| Trial number | 0.072 | 0.027 | 0.019 – 0.126 | 2.667 | 0.0077 |
| Pitch x Entropy (linear) | -0.186 | 0.081 | -0.344 – -0.027 | -2.300 | 0.0215 |
| Pitch x Entropy (quadratic) | 0.044 | 0.054 | -0.063 – 0.150 | 0.805 | 0.4207 |
| Pitch x Baseline Entropy | -0.036 | 0.081 | -0.194 – 0.122 | -0.443 | 0.6580 |
| Visual entropy (linear) x Baseline Entropy | 0.065 | 0.029 | 0.009 – 0.121 | 2.258 | 0.0240 |
| Visual Entropy (quadratic) x Baseline Entropy | -0.010 | 0.014 | -0.037 – 0.018 | -0.700 | 0.4838 |
| **Pitch x Pupil size (linear)** | **-0.249** | **0.069** | **-0.384 – -0.114** | **-3.615** | **0.0003** |
| **Pitch x Pupil size(quadratic)** | **-0.155** | **0.035** | **-0.224 – -0.086** | **-4.396** | **<0.001** |
| Pitch x Visual Entropy (linear) x Baseline Entropy | -0.135 | 0.067 | -0.266 – -0.004 | -2.027 | 0.0427 |
| Pitch x Visual Entropy (quadratic) x Baseline Entropy | 0.052 | 0.026 | 0.002 – 0.102 | 2.030 | 0.0424 |
| **Random Effects** | | | | | |
| $\sigma^2$ | 3.29 | | | | |
| $\tau_{00\ id}$ | 0.48 | | | | |
| $\tau_{11\ id.semitones}$ | 0.86 | | | | |
| $\rho_{01\ id}$ | -0.32 | | | | |
| Observations | 9831 | | | | |
| Marginal $R^2$ / Conditional $R^2$ | 0.544 / 0.633 | | | | |

**Table S12: Brain-behavior model predicting response speed**

| | RS | | | | |
|---|---|---|---|---|---|
| *Predictors* | *Estimates* | *std. Error* | *CI* | *t-value* | *p* |
| Intercept | 1.632 | 0.040 | 1.566 – 1.698 | 40.584 | <0.001 |
| Task ease | -0.103 | 0.004 | -0.109 – -0.096 | -25.930 | <0.001 |
| Entropy (linear) | -0.007 | 0.004 | -0.014 – 0.001 | -1.511 | 0.1308 |
| **Entropy (quadratic)** | **-0.012** | **0.004** | **-0.018 – -0.005** | **-3.079** | **0.0021** |
| Baseline Entropy | 0.003 | 0.005 | -0.006 – 0.012 | 0.545 | 0.5859 |
| Pupil size (linear) | -0.007 | 0.004 | -0.015 – -0.000 | -1.722 | 0.0850 |
| Pupil size (quadratic) | -0.004 | 0.003 | -0.009 – 0.000 | -1.641 | 0.1007 |
| Entropy (linear) x Entropy baseline | 0.001 | 0.004 | -0.007 – 0.008 | 0.134 | 0.8930 |
| Entropy (quadratic) x Entropy baseline | -0.003 | 0.003 | -0.008 – 0.002 | -1.088 | 0.2766 |
| **Random Effects** | | | | | |
| $\sigma^2$ | 0.16 | | | | |
| $\tau_{00\ id}$ | 0.04 | | | | |
| Observations | 9655 | | | | |
| Marginal $R^2$ / Conditional $R^2$ | 0.056 / 0.228 | | | | |
